# Identification of In Vivo Internalizing Cardiac-Specific RNA Aptamers

**DOI:** 10.1101/2024.08.13.607054

**Authors:** Chandan Narayan, Li-Hsien Lin, Maya N. Barros, Trent C. Gilbert, Caroline R. Brown, Dominic Reddin, Barry London, Yani Chen, Mary E. Wilson, Jennifer Streeter, William H. Thiel

## Abstract

**Background:** The pursuit of selective therapeutic delivery to target tissue types represents a key goal in the treatment of a range of adverse health issues, including diseases afflicting the heart. The development of new cardiac-specific ligands is a crucial step towards effectively targeting therapeutics to the heart.

**Methods:** Utilizing an *ex vivo* and *in vivo* SELEX approaches, we enriched a library of 2’-fluoro modified aptamers for ventricular cardiomyocyte specificity. Lead candidates were identified from this library, and their binding and internalization into cardiomyocytes was evaluated in both *ex vivo* and *in vivo* mouse studies.

**Results:** The *ex vivo* and *in vivo* SELEX processes generated an aptamer library with significant cardiac specificity over non-cardiac tissues such as liver and skeletal muscle. Our lead candidate aptamer from this library, CA1, demonstrates selective *in vivo* targeting and delivery of a fluorophore cargo to ventricular cardiomyocytes within the murine heart, while minimizing off-target localization to non-cardiac tissues, including the liver. By employing a novel RNase-based assay to evaluate aptamer interactions with cardiomyocytes, we discovered that CA1 predominantly internalizes into ventricular cardiomyocytes; conversely, another candidate CA41 primarily binds to the cardiomyocyte cell surface.

**Conclusions:** These findings suggest that CA1 and CA41 have the potential to be promising candidates for targeted drug delivery and imaging applications in cardiac diseases.

## INTRODUCTION

Cardiovascular disease (CVD) remains the leading cause of death in the US^1^ and is predicted to grow in prevalence over the next decade^2,3^. Despite this growing problem, the number of cardiovascular drugs entering the clinical pipeline has declined over the past twenty years ^4,5^. This gap in cardiovascular therapies has led to a call-to-action by the American Heart Association to “invest in the development of new approaches for the discovery, rigorous assessment, and implementation of new therapies”^6^. Although multiple pathways and therapeutic targets have been identified that could be modulated to treat diseases affecting the myocardium, a critical challenge in advancing cardiovascular therapeutics is the inability to *deliver* therapies that modulate these pathways specifically and efficiently to cardiomyocytes ^7,8^. Therefore, the identification of cardiotropic targeting ligands that can be administered systemically and accumulate within the heart is paramount to advance new therapies to treat CVD.

Among the most successful tissue-targeting strategies is the use of the GalNAC (triantennary N-acetyl galactosamine) moiety, which delivers cargo specifically to the liver by binding to the liver-specific asialoglycoprotein receptor. GalNAC-siRNA conjugates represent a definitive platform for delivering therapeutic siRNA cargos to the liver, culminating in the approval of four FDA-approved liver-targeted siRNA therapeutics^9^. Notably, each of these GalNAC-siRNA conjugates treats a different hepatic disease by targeting specific genes for silencing. The success of GalNAC as a targeting ligand underscores the significance of research aimed at discovering optimal tissue-tropic targeting ligands that enable entry into specific cell types while minimizing uptake by non-target tissues. Several tissue-targeting strategies have been evaluated for their ability to target the heart *in vivo*, including peptides, antibodies, and aptamers^10–16^. Each of these targeting ligands exhibits varying degrees of cardiac specificity over other tissue types in mice *in vivo*. However, few if any cardiac-specific targeting ligands have demonstrated significant specificity for cardiac tissues over the liver, and the capacity to internalize into cardiomyocytes rather than just bind to the cell surface.

Our objective in this study was to adapt aptamer technology to identify RNA aptamer ligands that preferentially target the heart and evaluate the ability of these aptamers to be internalized into adult ventricular cardiomyocytes within intact mouse cardiac tissue. Aptamers are short synthetic RNA or DNA oligonucleotides that are analogous to antibodies recognizing and binding to target epitopes with similar specificity and affinity as antibody-antigen interactions. Aptamers are a growing platform for diagnostics^17^, imaging^18^, therapeutics^19^, and targeted drug delivery^19,20^. Several aptamers are being evaluated clinically as diagnostic agents and therapeutics^21^, with two FDA-approved aptamers targeting VEGF^19^ and complement C5 for the treatment of macular degeneration^22^. Aptamers are identified using a process termed Systematic Evolution of Ligands by EXponential enrichment (SELEX)^23,24^. The SELEX process involves multiple rounds of selection to enrich a complex starting library, containing10^12^-10^36^ aptamer sequences toward those aptamers specific for a target of interest. SELEX selections for specific cell types, as opposed to a specific cell surface protein, have the advantage of being an unbiased approach for enriching cell-specific aptamers without prior knowledge of the proteins the aptamers may bind. We developed a combined *ex vivo* and *in vivo* SELEX process in mice and identified multiple cardiac-specific aptamers that can intenalize into cardiomyocytes.

## METHODS

### Aptamer RNA

Aptamer library or single aptamer template oligos (**Supplemental Table 1**) were chemically synthesized (IDT) as ssDNA with the first two 5’ nucleotides OMe-modified. The ssDNA templates were extended to dsDNA and the dsDNA was *in vitro* transcribed as 2’-fluoro-pyrimidine-modified aptamer RNA using the Y693F T7 RNA polymerase by previously published methods^25^. Fluorescently labeled aptamer RNA were chemically synthesized (Trilink) with a 12-carbon linker and Alexa647 fluorophore off the 5’ end of the aptamer. Prior to experiments, aptamer RNA was folded in binding buffer (200 mM HEPES pH 7.4, 500 mM NaCl, 20 mM CalCl_2_, 0.1% BSA) at 10 - 33.3 uM by heating to 95°C for 5 minutes followed by slow cooling to room temperature for 45 minutes and ice for at least 2 minutes. For *ex vivo* Langendorff heart perfusion experiments, folded aptamer RNA was diluted to the final experimental concentration in perfusion buffer. For *in vivo* mouse experiments, mice were immobilized and injected by tail vein with 4 nmoles of aptamer RNA at 33.3 uM in binding buffer.

### *Ex vivo* and *in vivo* SELEX

The *ex vivo* SELEX selection rounds involved perfusing a mouse heart with the aptamer library using a Langendorff heart preparation followed by isolation and purification of the ventricular cardiomyocytes. To minimize a sex bias or a single heart imparting a significant selection bias to the aptamer library, we used at least two mice per selection round while alternating between male and female mice during the SELEX process. For *ex vivo* SELEX selection rounds either male or female C57/bl6j (16-20 weeks of age) mouse hearts were perfused with a 37°C oxygenated wash perfusion buffer (123 mM NaCl, 4.7 mM KCl, 10 mM HEPES, 12 mM NaHCO_3_, 10 mM KHCO_3_, 0.6 mM KH_2_PO_4_, 0.6 mM Na_2_HPO_4_ 7H_2_O, 1.2 mM MgSO_4_) with 10 U/mL heparin (1,000 U/mL, NDC 63739-931-14) and 10 mg/mL yeast tRNA (ThermoFisher, AM7119) under constant pressure to remove blood. The mouse hearts were then perfused with a 37°C oxygenated aptamer library (75 – 300 nM) perfusion solution for 30 – 60 minutes. For *in vivo* SELEX selection rounds mice were injected with 4 nmoles aptamer library by tail vein. Following either aptamer library perfusion for *ex vivo* selection rounds or one-hour post-tail vein injection for *in vivo* selection rounds, the cardiomyocytes were dissociated and purified from non-cardiomyocytes by methods described below. Purified cardiomyocytes were pelleted by 1,500 xg centrifugation and the cardiomyocyte pellet resuspended in ∼1×10^5^ cells/mL TRIzol (Invitrogen, 15596026) containing 200 ug/mL Glycogen (Invitrogen, AM9515) and lysed using QIAshredder columns (Qiagen, 79654). Aptamer RNA was recovered from the TRIzol by organic extraction using 5PRIME phase lock tubes per the manufacturers protocol. Recovered aptamer library was reverse transcribed using Superscript IV (Invitrogen, 2848933) with a Sel2 3’ O-methyl-modified (OMe) primer (IDT, 5’-[UC]GGGCGAGTCGTCTG-3’ [UC] = OMe modification). The aptamer library cDNA was amplified by PCR using Q5 DNA polymerase (NEB, M0491) using the OMe-modified Sel2 3’ primer and the Sel2 5’ primer (IDT, 5’-TAATACGACTCACTATAGGGAGGACGATGCGG-3’). PCR product was purified using Qiaprep 2.0 spin columns (Qiagen, 27115) and the aptamer library dsDNA was used for *in vitro* transcription to produce aptamer RNA library for the next selection round.

### Cardiomyocyte dissociation and purification

Cardiomyocytes were dissociated by Langendorff heart perfusion of a 37°C digestion perfusion solution containing 300 U/mL collagenase (Worthington, LS004176). Collagenase digested tissue was minced and dissociated cardiomyocytes filtered through a 100 µm mesh filter (Falcon, 352360) into a 50 mL conical. Dissociated cardiomyocytes were purified from non-cardiomyocytes by three washes that included pelleting by 20 xg centrifugation and resuspending the pellet with 10 mL dissociation perfusion buffer containing 0.1% BSA (RPI, A30075). To determine purity of the cardiomyocyte purification, samples of the dissociated cardiomyocytes and from each of the three washes were fixed with 4% paraformaldehyde and stained for markers of cardiomyocytes, endothelial cells, fibroblasts, and smooth muscle as described below.

### NGS and aptamer bioinformatics

Recovered aptamer RNA from each selection rounds were reverse transcribed, and PCR amplified using barcoded Illumina compatible primers. PCR product was gel extracted, purified and the concentration of the purified dsDNA was determined by Qbit dsDNA HS assay (FisherScientific, Q32854). Barcoded amplicons of the selection rounds were pooled by equal molar amount and quality of the pooled sample was determined by Agilent Bioanalyzer. The pooled aptamer library amplicon sample was then submitted to the University of Iowa, Iowa Institute for Human Genetics for NGS on an Illumina NovaSeq 6000. Raw read data were uploaded to Galaxy^26^ and processed into a non-redundant database comprised of variable region sequence information and read counts^27,28^. The non-redundant database was filtered based on normalized aptamer abundance (read counts) and persistence (number of rounds an aptamer was detected within)^29^. The filtered non-redundant database was converted into a FASTA formatted file containing the full-length aptamer RNA sequences, which were then clustered for sequence similarity (edit distance) and structure similarity (tree distance) using AptamRunner^30^. Clustering results were visualized within Cytoscape^31^ with log2 fold enrichment data and read count data used to determine node color and size respectively. Candidate aptamers were identified from separate sequence/structure families that exhibited a positive log2 fold enrichment. Aptamer tertiary structures were predicted using trRosettaRNA^32^ and visualized using Mol*^33^.

### *Ex vivo* mouse heart perfusion

Mouse hearts were perfused as described for the *ex vivo* SELEX methods with 37°C oxygenated heparinized perfusion buffer under constant pressure as follows: wash ∼5 minutes and 150 nM aptamer (*in vitro* transcribed or Alex647 chemically synthesized) for 45 minutes. Experiments that isolated and purified the cardiomyocytes followed the aptamer prefusion with a cardiomyocyte dissociation and purification as described above. Experiments that evaluated aptamer binding versus internalization treated half of the cardiomyocytes with 5,000 U/mL RNase T1 (ThermoScientific, FEREN0541) and 2,000 gel units/mL micrococcal nuclease (NEB, M0247S) for ten minutes on ice during the second wash step. Following the final wash cardiomyocytes were counted and aliquoted into 1×10^5^ cells per sample for TRIzol extraction and quantification using methods described below. *Ex vivo* experiments that imaged the perfused hearts followed the aptamer solution perfusion with a 15-minute wash with perfusion buffer followed by 4% paraformaldehyde for 10 minutes.

### *In vivo* mouse tail-vein injection

Mice were injected via the tail vein with 4 nmoles of aptamer as described for the *in vivo* SELEX methods (*in vitro* transcribed or Alex647 chemically synthesized). Experiments evaluating aptamer binding and internalization removed the heart one-hour post-injection to isolate and treat cardiomyocytes as described with *ex vivo* perfused mouse hearts to assess aptamer binding versus internalization. Experiments evaluating aptamer tissue localization recovered cardiac and non-cardiac tissue and either fixed tissue using 4% paraformaldehyde or recovered aptamer from ∼25 – 30 mg tissue using 1 mL TRIzol per 50 mg tissue. Fixed tissue was immune-stained and imaged by confocal microscopy as described below. Cardiac and non-cardiac tissue was lysed for TRIzol extraction and quantification by methods described below.

### Aptamer TRIzol extraction and Reverse Transcription quantitative PCR (RT-qPCR)

Aptamer RNA was recovered by TRIzol extraction from either isolated cardiomyocytes (4×10^5^ cells/mL TRIzol), or mouse tissue (50 mg/mL TRIzol). When possible, those performing the TRIzol extraction and RT-qPCR were blinded to the aptamer treatment. Isolated cardiomyocytes were lysed in TRIzol using QIAshredder spin columns. Tissue from mice was lysed in TRIzol using reinforced 2 mL homogenizer tubes with 2.8-mm ceramic beads (Bertin, CK28R) homogenized by a Precellys 24 Homogenizer at 6,500 rpm 3x 15s cycles with 15s between cycles. For all samples TRIzol was supplemented with 200 ug/mL glycogen (Invitrogen, AM9515). Organic extraction followed the manufacturers protocol using 5PRIME phase lock tubes to facilitate collection of the aqueous phase. The aqueous phase was treated with RNase A (Thermo Scientific, 2766319) to degrade endogenous RNA, but not the 2’-fluoro modified aptamer RNA. Isopropanol precipitated aptamer RNA was washed with 75% ethanol, air dried and resuspended in 200 uL PCR-grade H_2_O per mL TRIzol used for lysis. TRIzol recovered aptamer RNA and equal volume aptamer RNA standards (1:5 dilution from 1 nM aptamer RNA) were reverse transcribed using Superscript IV and quantified by using the QuantStudio 3 Real-Time PCR system with iQ SYBR Green Supermix (Bio Rad). RT-qPCR data were analyzed using Applied Biosystems Design and Analysis Software 2.6.0.

### Immuno-staining and confocal microscopy

Procedures similar to those described in previous publications^34–37^ were used for multiple immunofluorescent staining of isolated cells and tissue. Cells or tissue sections were mounted on Colorfrost Plus microscope slides (Fisher Scientific), air-dried and washed with PBS (phosphate buffered saline), they were than blocked with 10% donkey normal serum (Jackson Immuno Research Lab., USA), and then incubated with primary antibodies (see **Supplemental Table 2** for sources and dilution of each antibody) in 10% donkey normal serum at 25°C overnight. They were then incubated with appropriate affinity purified fluorescent dye-conjugated secondary antibodies (Alexa Fluor 488 or Alexa Fluor 568 or Alexa Fluor 647 conjugated, all at 1:200 dilution, all from Jackson Immuno Research Lab.) at 4°C overnight after thoroughly washed with PBS. The slides were then washed and stained with a fluorescent nucleus dye (TOPRO-3,1: 2000 dilution, Molecular Probes or SYTOX green, 1:5000 dilution, Invitrogen), and/or a fluorescent actin dye (Alexa Fluor 568-Phalloidin or Alexa Fluor 488-Phalloidin, both at 1:40 dilution, Molecular Probes) for 15 min. We then washed the slides and cover-slipped them with Prolong Diamond Antifade Reagents (Invitrogen-Molecular Probes, USA). Multiple-label immunofluorescent staining was performed with primary antibodies that were raised in different species. We analyzed stained cells or tissue sections with a Zeiss LSM 710 confocal laser-scanning microscope as described in earlier publications^34–36^. Digital confocal images were obtained and processed with software provided with the Zeiss LSM 710.

### Data analysis and statistics

Data are presented as mean ±SEM with sample sizes of n = 3 - 10. Statistical tests include 2-way ANOVA with multiple comparisons and Ordinary 1-way ANOVA with multiple comparisons. A *P* value of < 0.05 considered significant. Data analysis and statistics were conducted using Microsoft Excel and GraphPad Prism 10.

## RESULTS

### *Ex vivo* and *in vivo* SELEX for cardiac-specific RNA aptamers

Aptamers specific for ventricular cardiomyocytes were identified using a SELEX process that incorporated a combination of *ex vivo* and *in vivo* selection rounds (**Figure 1A**). To ensure the enrichment of cardiac-specific aptamers during SELEX, we critically evaluated cardiomyocyte isolation and purification methods aimed at retaining cardiomyocytes while eliminating non-cardiac cell types. Our finding revealed that a series of three washes did not significantly impact on the cardiomyocyte fraction, but each wash resulted in a marked decline in non-cardiomyocyte cells, including endothelial cells, fibroblasts, and smooth muscle cells (**Supplemental Figure 1**).

**Figure 1:**
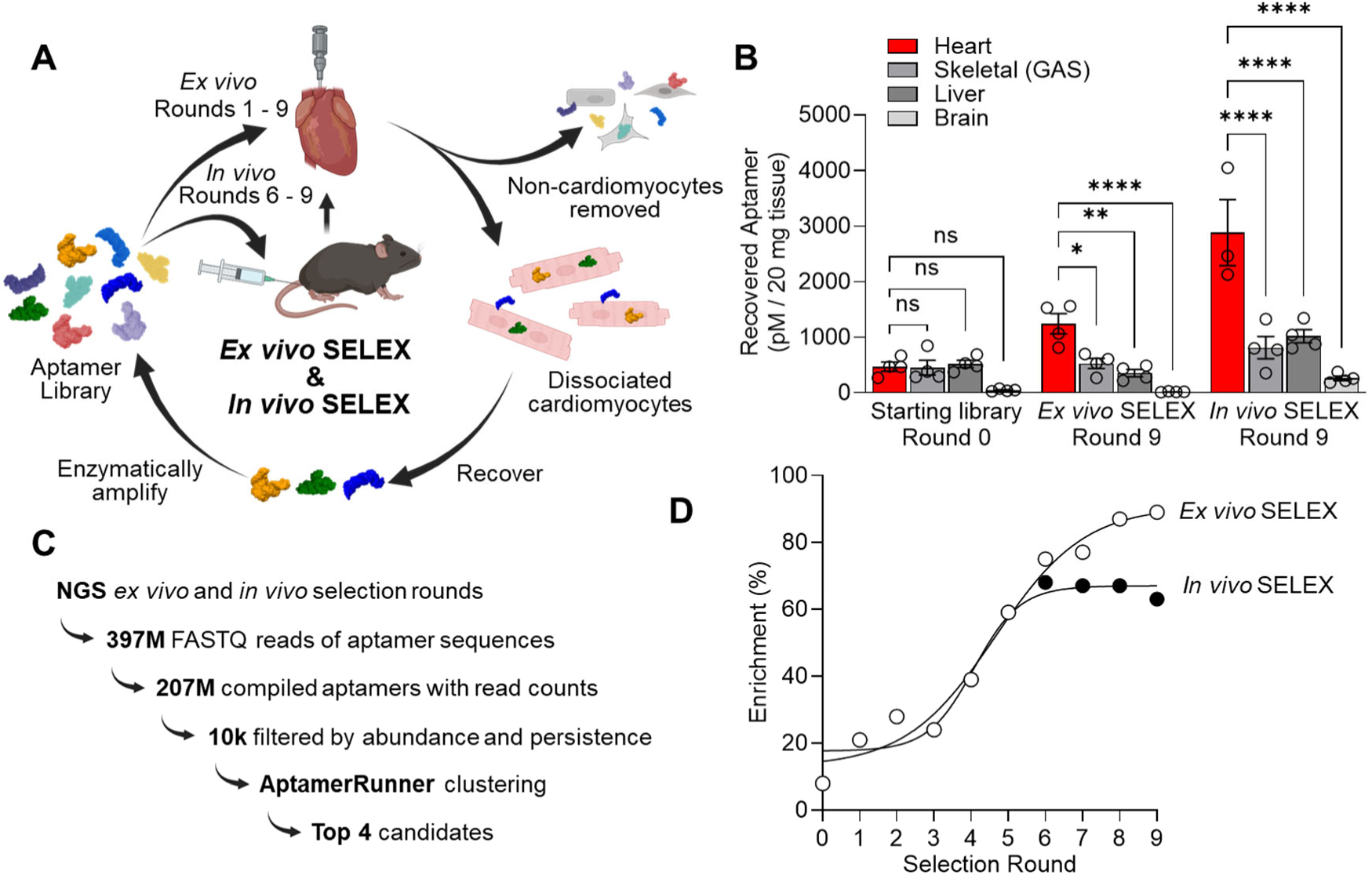
*Ex vivo* and *in vivo* cardiac SELEX. **A)** Schematic of the *ex vivo* (rounds 1 – 9) and *in vivo* (rounds 6 – 9) SELEX strategy. **B)** Cardiac tissue specificity of the *ex vivo* final selection round 9 and final *in vivo* selection round 9 as compared to the starting aptamer library (round 0). 2-way ANOVA with multiple comparisons; * p<0.05, ** p<0.01, ****p<0.0001, ns = not significant, n= 3 - 4. **C)** Schematic of NGS data acquisition following SELEX, and analysis strategy of the NGS data to identify lead cardiac aptamer candidates. **D)** Sequence enrichment as determined by the NGS data for the *ex vivo* cardiac SELEX selection rounds and *in vivo* cardiac SELEX selection rounds.

Using this cardiomyocyte purification protocol, we conducted five rounds of *ex vivo* SELEX with mouse hearts. *Ex vivo* selection rounds involved retrograde perfusion of the mouse heart with the aptamer library for one hour, accompanied by the isolation of cardiomyocytes. Following these five rounds, we proceeded with four *in vivo* selection rounds in parallel with an additional four *ex vivo* selection rounds. The *in vivo* selection rounds involved injecting the aptamer library by tail-vein into a mouse, waiting one hour and then isolating the cardiomyocytes from the mouse hearts. Integrating *in vivo* selection rounds along with *ex vivo* selection rounds enabled a direct comparison between a SELEX process conducted entirely *ex vivo* and a SELEX process that combined both *ex vivo* and *in vivo* approaches.

We evaluated the final selection rounds of the *ex vivo* only cardiac selection and *ex vivo* & *in vivo* combined selection for tissue selectivity (**Figure 1B**) within the heart, skeletal muscle (Gastrocnemius), the liver, and the brain. Skeletal muscle and the liver were chosen because they have been noted as the largest off-target tissue reservoirs for other cardiac targeting molecules ^15,38,39^. Brain tissue was used as a negative control, as aptamers are unlikely to cross the blood-brain barrier^40^. Both *ex vivo* and *in vivo* aptamer libraries were observed to localize primarily within the heart, with significantly lesser amounts observed in skeletal muscle and liver. As expected, brain tissue exhibited the least amount of aptamer localization. Importantly, aptamers from the final *in vivo* selection round demonstrated greater specificity for heart over skeletal muscle and liver compared to the final *ex vivo* only selection round. These data suggest that while *ex vivo* selection pressure was adequate to generate cardiac-specific aptamers, applying *in vivo* selection pressure significantly enhance the tissue specificity of the aptamer library.

### Next-generation sequencing (NGS) and bioinformatics analysis

To identify the aptamers enriched during SELEX, we prepared the starting aptamer library and all subsequent selection rounds for next-generation sequencing (NGS). NGS identified 397,642,327 aptamer sequences reads, representing 207,6024,599 unique aptamer sequences (**Figure 1C**). From these NGS data, we examined the ratio of total aptamer sequence reads (Total) to the number of unique aptamers sequences (Unique) attained from each selection round to determine the degree of the aptamer library enrichment (Selection Round Enrichment % = 1-[Unique/Total]). For the *ex vivo* only selection rounds and the combined *ex vivo* & *in vivo* selections, the NGS indicates 50% enrichment was achieved between rounds three and five, with maximum enrichment of the aptamer library achieved after round six (**Figure 1D**). Interestingly, we observed greater enrichment with the *ex vivo* only selection as compared to the combined *ex vivo* & *in vivo* selection. This may be due to unanticipated differences between *ex vivo* selection conditions and *in vivo* selection conditions, where a degree of library complexity is retained despite an increase in library specificity for cardiac tissue with the *in vivo* selection rounds.

All sequenced aptamers were compiled into a non-redundant database that tracked read counts of each unique aptamer sequence across all selection rounds. This database was filtered based on an aptamer persistence and abundance analysis of the selection rounds, as compared to the starting aptamer library, to include 10,657 aptamers with at least 135 reads and found within at least four selection rounds. These 10,657 aptamers were clustered for sequence and predicted structure similarity using our AptamerRunner clustering algorithm (**Supplemental Figure 2**). Within the sequence and structure clusters, aptamers were ranked by log2 fold round-to-round enrichment. From different clusters, we identified four cardiac aptamer (CA) candidates - CA1, CA3, CA12 and CA41 - for experimental validation. A negative control aptamer was identified as a sequence observed within the starting aptamer library but absent in any sequenced selection rounds.

### Validation of candidate aptamer cardiomyocyte affinity and assessment of aptamer localization to cell surface versus cellular internalization

We aimed to quantify the fraction of the lead cardiac aptamers that either bound to the surface of cardiomyocytes or were internalized within them. Understanding whether an aptamer binds to the surface or is internalized into cardiomyocytes will aid in determining potential applications, such as delivering nanoparticles to the cell surface or delivering siRNA within the cell. To quantify the fraction of an aptamer internalized into cardiomyocytes, we developed a novel internalization assay that employs a cocktail of bacterial RNases that degrade 2’-fluoro-modified aptamers (**Supplemental Figure 3**). This assay specifically degrades all bound aptamers present on the cell surface, while leaving those internalized into cardiomyocytes unaffected (**Figure 2A**). By utilizing this assay, we are able to measure both the bound and internalized fractions of aptamers in RNase untreated cells, as well as the internalized fraction in RNase cocktail-treated cells. From these measurements, the ratio of internalized versus bound aptamers can be accurately calculated.

**Figure 2:**
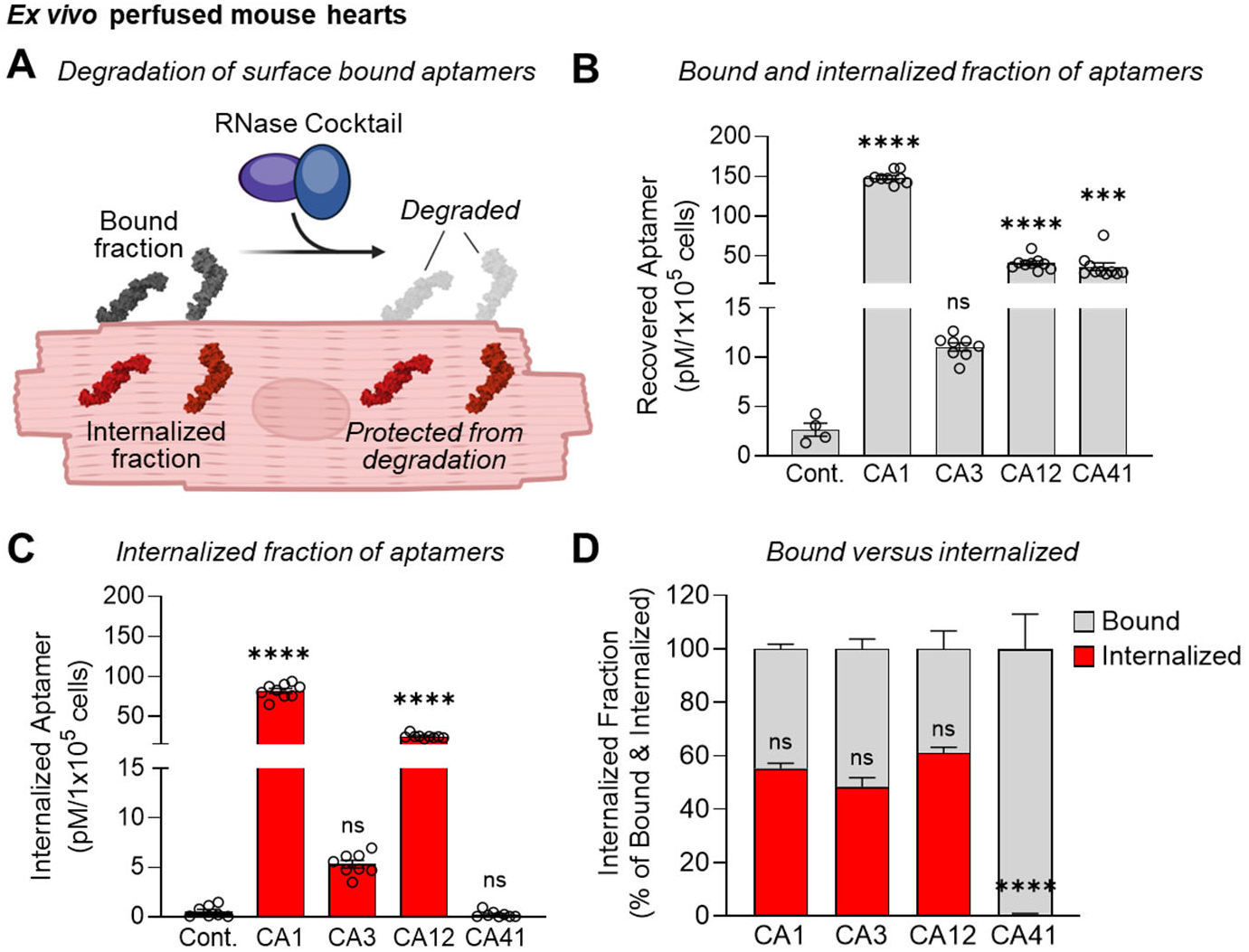
Quantifying bound versus internalized fraction of cardiac aptamers. **A)** Schematic of a novel internalization assay to quantify the amount of internalized aptamer by using a cocktail of bacterial RNases to degrade bound aptamers. **B)** Bound and internalized fraction (untreated) of aptamers, and **C)** internalized fraction (RNase cocktail treated) of aptamers associated with purified ventricular cardiomyocytes isolated from mouse hearts perfused *ex vivo*. Ordinary 1-way ANOVA with multiple comparisons against control; *** < 0.001, **** < 0.0001, ns = not significant, n = 4 - 10. **D)** Calculated fraction of bound aptamer versus internalized. 2-way ANOVA with multiple comparisons between all RNase cocktail treated samples; **** < 0.0001.

Each lead candidate cardiac aptamer and the control aptamer were perfused *ex vivo* into mouse hearts. Ventricular cardiomyocytes were then isolated from the *ex vivo* perfused mouse hearts, and the purified cardiomyocytes were either untreated (**Figure 2B**) or treated with the RNAase cocktail (**Figure 2C**). For the untreated cardiomyocytes, which constitutes the bound and internalized fraction of aptamer, we observed that candidate cardiac aptamers CA1, CA12, and CA41, but not CA3, were associated significantly more with the cardiomyocytes as compared to the negative control aptamer. The RNase cocktail-treated cardiomyocytes indicate that significantly more CA1 and CA12 internalized into the cardiomyocytes than the control aptamer. Whereas the CA3 and CA41 cardiac aptamers was observed to not have a significant internalized fraction as compared to the control aptamer. The CA1 aptamer was observed to have the greatest amount of aptamer associated with both the untreated and RNase cocktail-treated cardiomyocytes. The CA3 aptamer exhibited a trend towards targeting cardiomyocytes, but the observed effect was not found to be statistically significant. Data from RNase cocktail-treated and untreated cardiomyocytes indicate that 55% of CA1, CA3, and CA12 internalize into the cardiomyocytes with no significant difference between the fraction internalized for these cardiac aptamers (**Figure 2D**). From these data we conclude that CA1 would be the ideal internalizing cardiac aptamers, and that CA41 would be the ideal aptamer that binds to the cell surface but does not internalize into cardiomyocytes.

To investigate the potential of the cardiac aptamers to deliver a small molecule to ventricular cardiomyocytes and to verify our *ex vivo* aptamer binding and internalization results, we modified these cardiac aptamers to carry a fluorescent tag. An Alexa 647 fluorophore (AF647) was appended to the 5’ end of CA1, CA41, and the control aptamer via a 12-carbon linker (**Figure 3A**). We perfused these fluorescent-tagged cardiac aptamers ex vivo into mouse hearts, followed by sectioning, and imaging by confocal microscopy for aptamer fluorescence within the left ventricle (**Figure 3B**) and right ventricle (**Supplemental Figure 4**). We observed significantly more CA1-AF647 and CA41-AF647 fluorescence throughout the left and right ventricle wall as compared to control-AF647.

**Figure 3:**
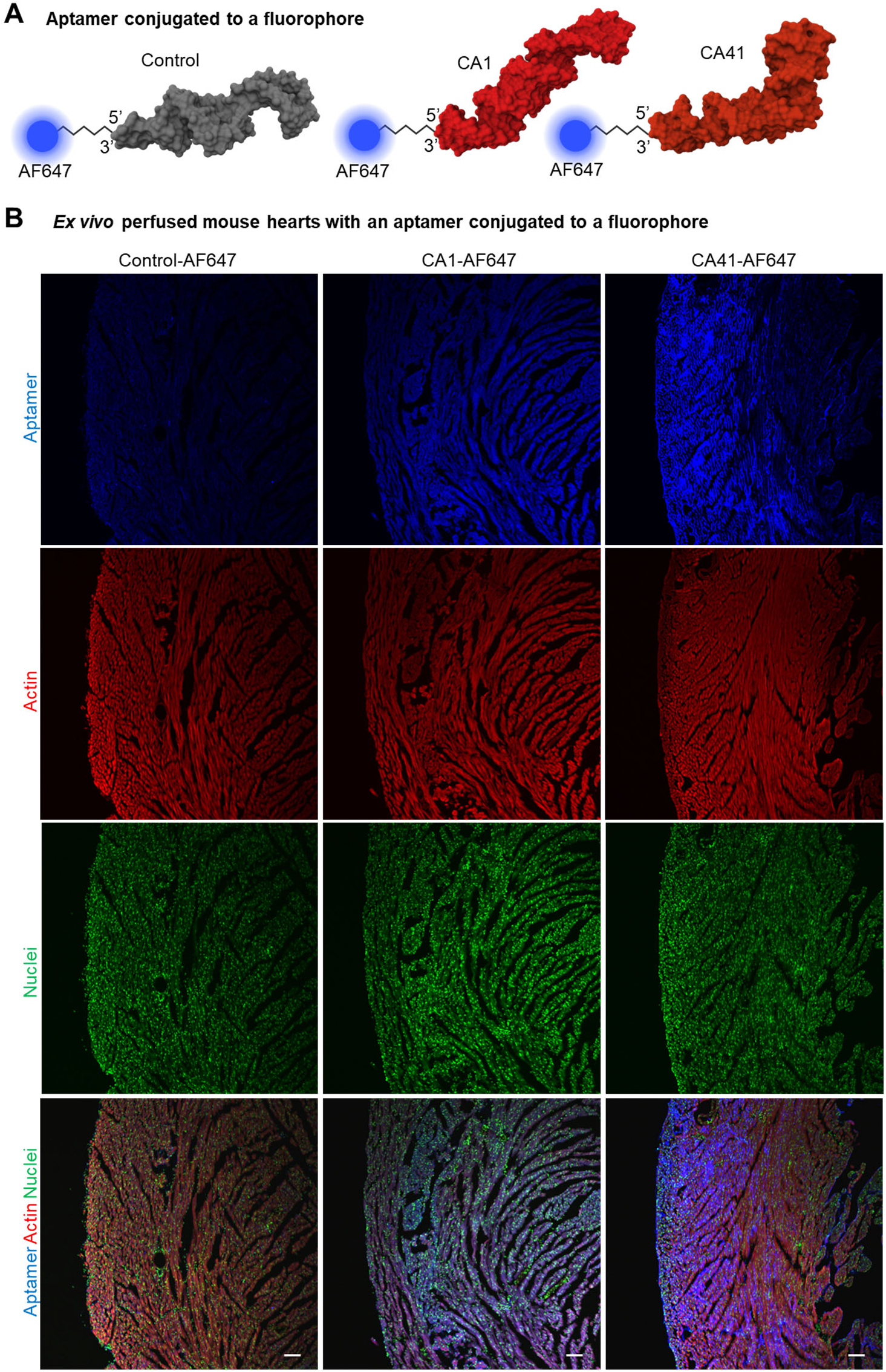
Cardiac aptamer conjugated with an Alexa647 (AF647) fluorophore. **A)** Tertiary structure prediction of the CA1-AF647, CA41-AF647 and control-AF647 aptamer conjugated at the 5’ end with AF647 via a 12-carbon linker. **B)** Images of left ventricle sections from mouse hearts perfused *ex vivo* with AF647 conjugated aptamers (blue), and stained for actin (red) and nuclei (green). Scale = 100 µm

Closer examination of the left ventricular cardiomyocytes (**Figure 4A, top**) indicates that CA1-AF647 is bound and internalized into cardiomyocytes, whereas CA41-AF647 appears to only to bind to the cardiomyocytes. Limited binding or internalization of the control-AF647 was observed. These fluorescence patterns for CA1-AF647, CA41-AF647 and the control-AF647 were consistent throughout the left and right ventricles across the basal, mid and apex myocardium (**Supplemental Figure 5**). Importantly, imaging of vessels within the myocardium reveals that cardiac aptamers CA1-AF647 and CA41-AF647 do not interact with the endothelium or medial layer of arteries within the heart (**Figure 4A, bottom**).

**Figure 4:**
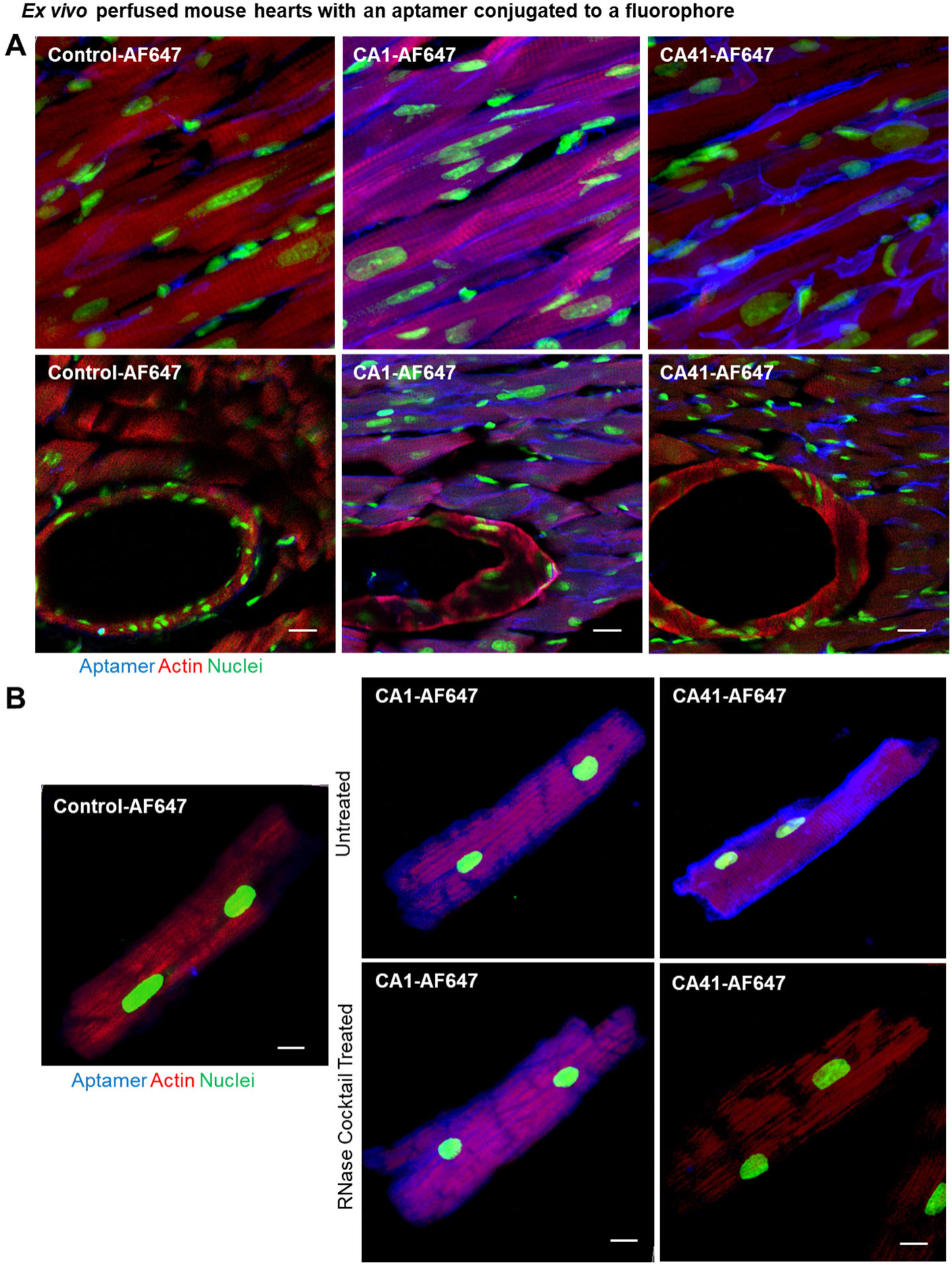
Cardiac aptamers conjugated to a fluorophore bind and internalize into left ventricular cardiomyocytes. **A)** Sections of left ventricular myocardium of mouse hearts perfused *ex vivo* with CA1-AF647, CA41-AF647, or control-AF647. **B)** Left ventricular cardiomyocytes from mouse hearts perfused *ex vivo* with CA1-AF647, CA41-AF647 or control-AF647 were treated with or without the RNase cocktail. Aptamer (blue), actin (red), and nuclei (green). Scale = 25 µm

To further visualize how these aptamers interact with the cardiomyocytes, we isolated and imaged the individual ventricular cardiomyocytes from mouse hearts perfused *ex vivo* with the fluorescently labeled aptamers (**Figure 4B**). We then treated a portion of the isolated cardiomyocytes with the RNase cocktail. We observed that CA1-AF647 predominantly localized to the interior of cardiomyocytes, while CA41-AF647 localized mainly to the cell surface (**Figure 4B, Untreated**). These observations were supported by the CA1-AF647 fluorescent signal being maintained and the CA41-AF647 signal was lost after RNAse cocktail treatment (**Figure 4B, RNAse Cocktail Treated**). These observations corroborate our previous results: CA1 internalizes into cardiomyocytes, while CA41 binds only to the cardiomyocyte cell surface. These data demonstrate that CA1 can deliver a small molecule into cardiomyocytes, while CA41 can target a small molecule to the cell surface of cardiomyocytes.

### *In vivo* cardiac aptamer tissue specificity

To ensure the cardiac aptamers could be administered systemically, travel through the circulatory system while bypassing off-target tissues, and efficiently localize to the heart before being renally cleared, we assessed their ability to perform these functions *in vivo* in mice. Specifically, we evaluated the cardiac aptamers CA1 and CA41 for their capacity to bind and internalize into cardiomyocytes one hour after injection via the tail vein (**Figure 5A**). CA1 demonstrated elevated recovery from isolated mouse cardiomyocytes post-injection, as compared to the control aptamer, further supporting its cardiac specificity. Treating these cardiomyocytes with an RNase cocktail did not substantially decrease the recovered CA1 amount, suggesting that most of CA1 resides within the cardiomyocytes. Conversely, CA41 did not bind cardiomyocytes significantly more than the control aptamer *in vivo*. However, after RNase cocktail treatment, significantly more CA41 was detected than the control aptamer. Yet, despite this significant difference, the amount of CA41 remained similar to the amount recovered from the untreated cardiomyocytes. These results suggest that a small fraction of CA41 internalizes *in vivo* and its binding to cardiomyocytes might be transient, potentially due to renal clearance of the aptamers.

**Figure 5:**
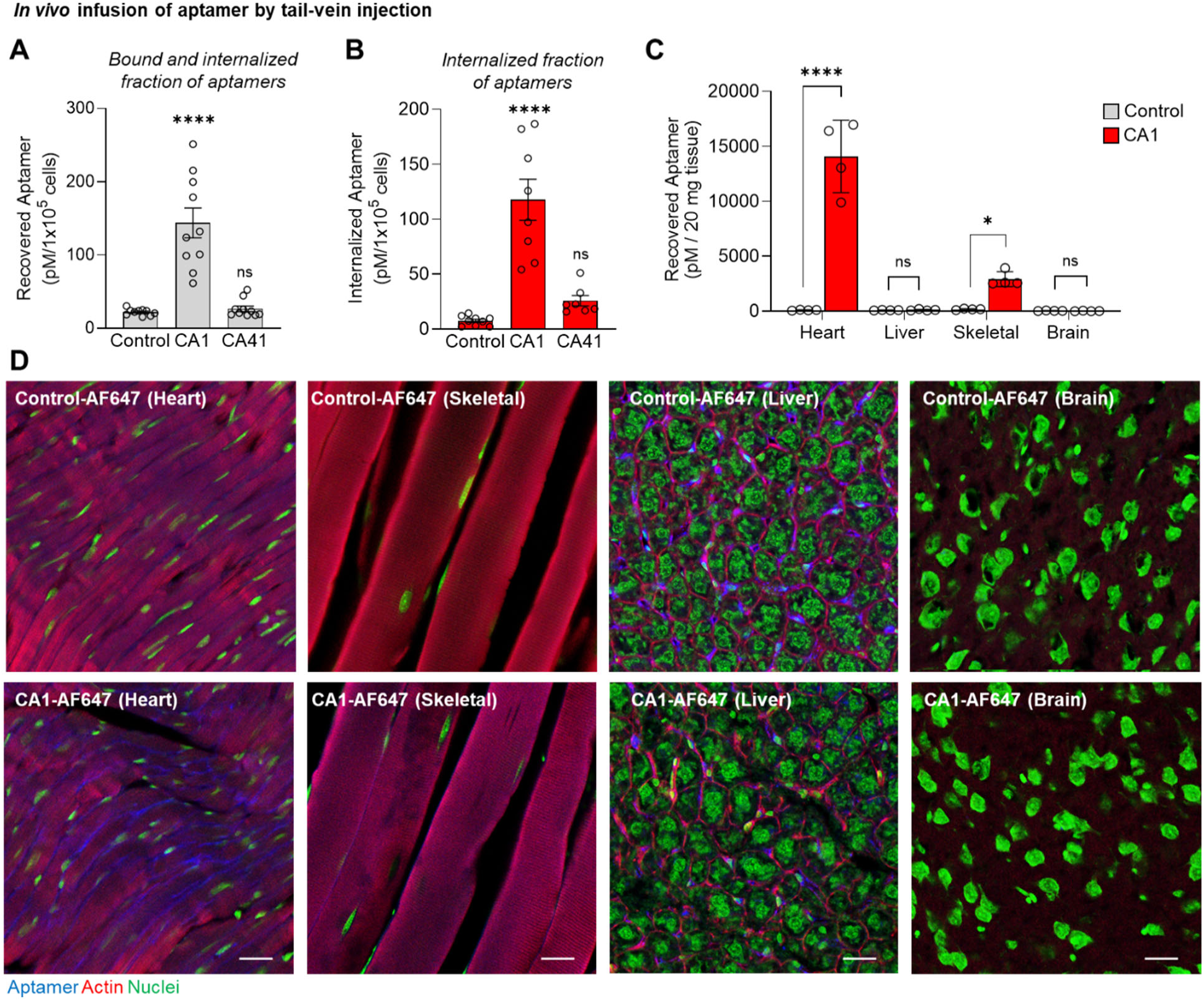
*In vivo* localization of cardiac aptamers. Purified cardiomyocytes either **A)** untreated or **B)** treated with RNase cocktail from mice injected via the tail vein with either CA1 or control aptamer. Ordinary 1-way ANOVA with multiple comparisons; **** < 0.0001, ns = not significant, n = 7 - 10. **C)** Quantification of CA1 and control aptamer *in vivo* localization to the heart, liver, skeletal muscle (gastrocnemius), and brain following injection into mice via the tail vein. 2-way ANOVA with multiple comparisons; * < 0.05, **** < 0.0001, ns = not significant, n = 4. **D)** *In vivo* localization of CA1-AF647 and control-AF647 after injection via the tail vein to the heart, liver, skeletal muscle (gastrocnemius), and brain. Aptamer (blue), actin (red), and nuclei (green). Scale = 25 µm

We next evaluated the tissue selectivity of CA1 and the control aptamer for the heart compared to non-cardiac tissues following injection via the tail vein. We observed significantly higher levels of CA1-AF647 aptamer in the heart than in skeletal muscle (gastrocnemius), liver, and brain (**Figure 5B**). These data support the conclusion that CA1 exhibits highly selective cardiac specificity *in vivo*. Although CA1 also showed significant targeting of skeletal muscle tissue compared to the control aptamer, its cardiac selectivity is 4.8-fold higher than that observed for skeletal muscle.

To determine whether CA1 can deliver cargo to cardiomyocytes *in vivo*, we injected fluorescent-tagged CA1-AF647 or control-AF647 into mice via the tail vein and imaged tissue sections one-hour post-injection. We observed that CA1-AF647 fluorescence was significantly higher in the heart compared to skeletal muscle (gastrocnemius), liver, and brain, as well as compared to control-AF647 within the heart (**Figure 5C**). More CA1-AF647 signal was observed within skeletal muscle than control-AF647, but the signal in the heart was 4.8 fold greater than that in skeletal muscle, which is consistent with the quantification of CA1 in heart and skeletal muscle by qPCR. More control-AF647 appeared localized to the liver than CA1-AF647, and no detectable amounts of either CA1-AF647 or control-AF647 were observed within the brain. These fluorescent data correspond to the quantitative PCR results of CA1 recovery from cardiac and non-cardiac tissues. Importantly, these results demonstrate that CA1 can be injected systemically, travel through the circulatory system, avoid localization to off-target tissues such as the liver, effectively and specifically localize to the heart, and deliver cargo into cardiomyocytes *in vivo*.

## CONCLUSIONS

We compared *ex vivo* only and combined *ex vivo* & *in vivo* SELEX processes to assess tissue selectivity of aptamers targeting the heart. Our findings revealed that while both methods generate aptamers primarily localizing in heart tissue, the combined approach enhances tissue specificity over skeletal muscle and liver. Significantly, aptamers from *in vivo* selection showed increased heart specificity, underscoring the value of incorporating *in vivo* selection rounds to improve the precise targeting of cardiac-specific molecules. Our data suggests that the *in vivo* selection rounds are crucial for imparting tissue specificity to the aptamer library. However, we believe that the *ex vivo* selection rounds present an opportunity to establish the initial specificity of the aptamer library under carefully controlled conditions, such as aptamer concentration and perfusion time. Importantly, both *ex vivo* and *in vivo* selection rounds included the isolation and purification of ventricular cardiomyocytes, which is necessary to avoid non-cardiomyocyte bias in the library and ensuring that only aptamers escaping the vasculature are enriched.

We observed distinctive interactions of CA1 and CA41 aptamers with cardiomyocytes *ex vivo*. Fluorescently labeled CA1-AF647 shows a significant internalization within the cells, retaining its fluorescent signal even after RNase treatment, indicating its potential for intracellular delivery. In contrast, CA41-AF647 primarily binds to the cell surface of cardiomyocytes and its signal diminishes post-RNase treatment, suggesting its use for surface-targeted delivery. CA1 demonstrates remarkable cardiac specificity when administered *in vivo*, as evidenced by significantly higher fluorescence in heart tissue compared to skeletal muscle, liver, and brain. Systemic injection of fluorescent-tagged CA1 shows that it selectively targets and delivers cargo to cardiomyocytes in the heart, avoiding significant off-target localization. These findings are supported by both fluorescent imaging and quantitative PCR results, underscoring CA1’s potential for targeted cardiac therapies.

We were able to define the binding versus internalizing capability of CA1 and CA41 by applying a novel RNase cocktail treatment. Understanding these characteristics of CA1 and CA41, to either internalize or bind cardiomyocytes, can be strategically utilized for targeted therapeutic interventions in cardiac cells. Surface-binding aptamers, such as CA41, can be engineered to potentially trigger extracellular signaling pathways or deliver nanoparticles that remain on the cell membrane, serving as targeted drug delivery systems for cardiac conditions. On the other hand, aptamers that are internalized into cardiomyocytes like CA1 open up more sophisticated therapeutic possibilities, such as the delivery of siRNAs, as accomplished by GalNac for targeting the liver. This intracellular delivery of siRNAs can silence specific genes, offering a precise and targeted approach for treating various cardiac diseases. Furthermore, understanding these interactions can help develop multifunctional aptamers that combine surface binding with intracellular delivery, providing a comprehensive tool for both diagnostic and therapeutic applications targeting the heart.

While the CA1 and CA41 cardiac-specific aptamers show promise in targeting cardiomyocytes, our data indicates clear areas for improving the cardiac SELEX process. Future iterations of cardiac SELEX could utilize human induced pluripotent stem cell cardiomyocytes to support the cross-reactivity of the aptamers with human cardiomyocytes. Additionally, cardiac-SELEX and other cell-based SELEX could employ RNases to degrade the bound fraction of aptamers, thereby driving enrichment toward those that best internalize into target cells. Together, these techniques could enhance aptamer enrichment, favoring those that most effectively internalize into human cardiomyocytes.

In summary, the continuous advancements in the cardiac SELEX methodology promise to revolutionize the landscape of cardiovascular therapeutics and diagnostics. By embracing innovative techniques, the SELEX process is poised to deliver unprecedented precision in targeting cardiomyocytes. This will ultimately enhance patient outcomes and drive forward our capabilities in managing and treating cardiac diseases.

## DATA AVAILABILITY

Source data will be provided by the corresponding authors upon reasonable request.

## ACKNOWLEDGMENTS

This work was supported by grants from: the National Institutes of Health to WHT (R01HL139581, R01HL157956), American Heart Association to WHT (18IPA34170406) and to JS (23IPA1054531), and the US Department of Veterans’ Affairs to MW (I01 BX001983, I01 BX000536). We would like to thank the Iowa City VA Medical Center for the use of the confocal microscope.

## SUPPLEMENTAL FIGURES AND TABLES

**Supplemental Figure 1:**
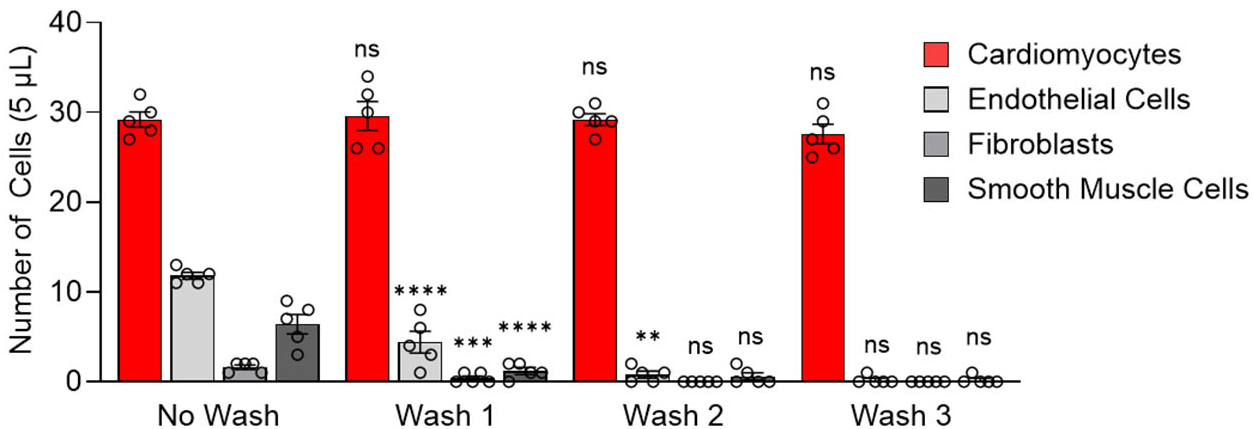
Purification of cardiomyocytes from non-cardiomyocytes cells following collagenase digestion of *ex vivo* perfused mouse hearts. 1-way ANOVA for each cell type with multiple comparisons between sequential washes; ** p<0.01, *** p<0.001, **** p<0.0001, ns = not significant, n = 5.

**Supplemental Figure 2:**
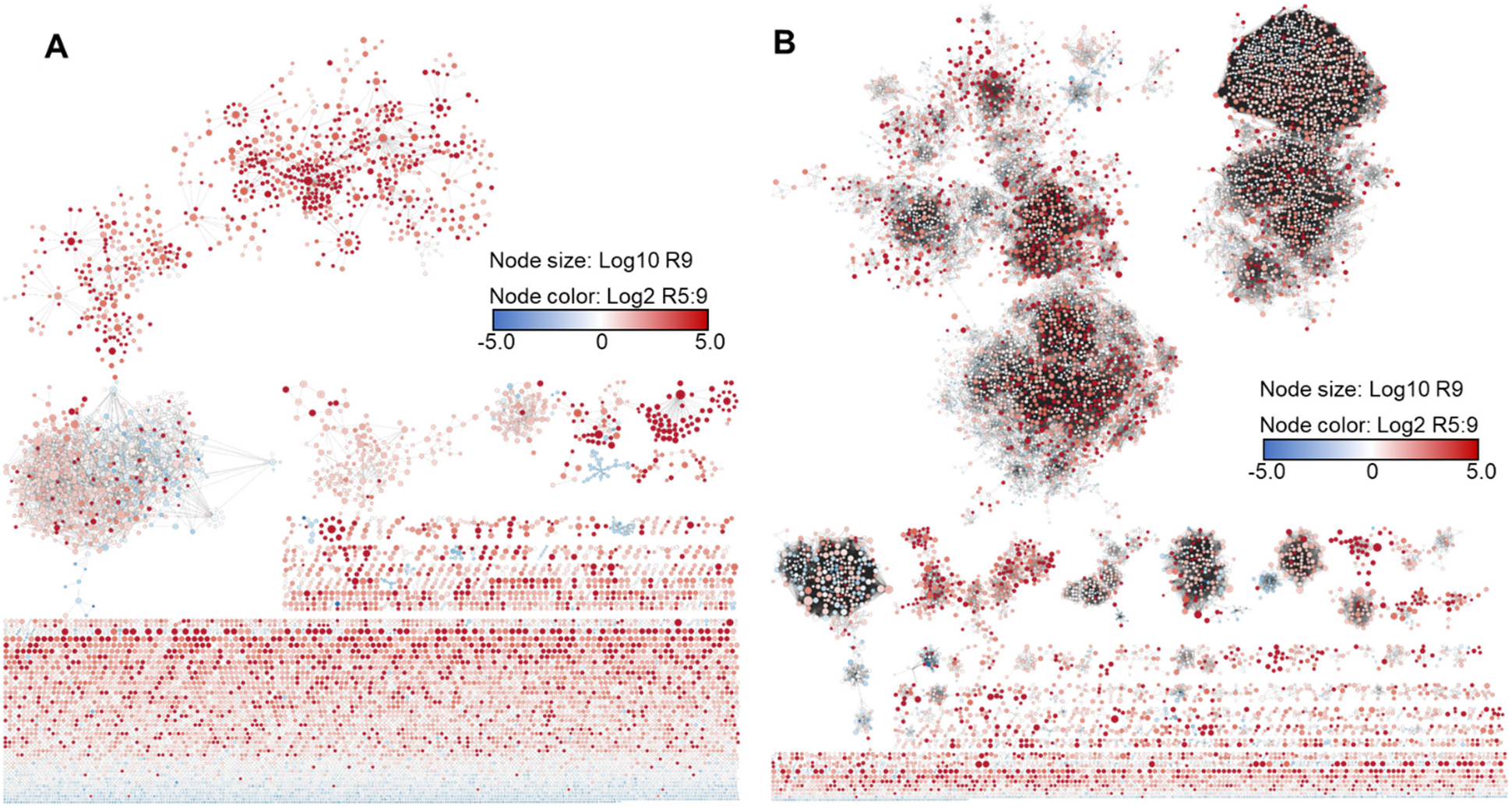
Clustering of cardiac aptamers by the AptamerRunner clustering algorithm. **A)** Cytoscape visualization of aptamer sequences related by sequence similarity (edit distance 1), and by **B)** structure similarity (tree distance 3).

**Supplemental Figure 3:**
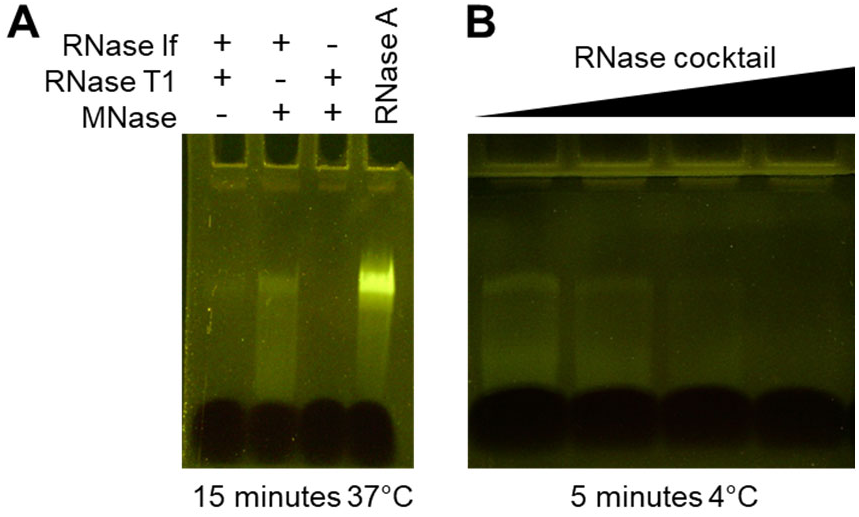
RNase cocktail used to degrade cell-surface bound aptamers. **A)** Evaluation of bacterial RNases and nuclease for their ability to degrade 2’-fluoro-pyrimidine-modified aptamers as compared to RNase A. **B)** Dose dependence of the RNase cocktail at 4°C for 5 minutes.

**Supplemental Figure 4:**
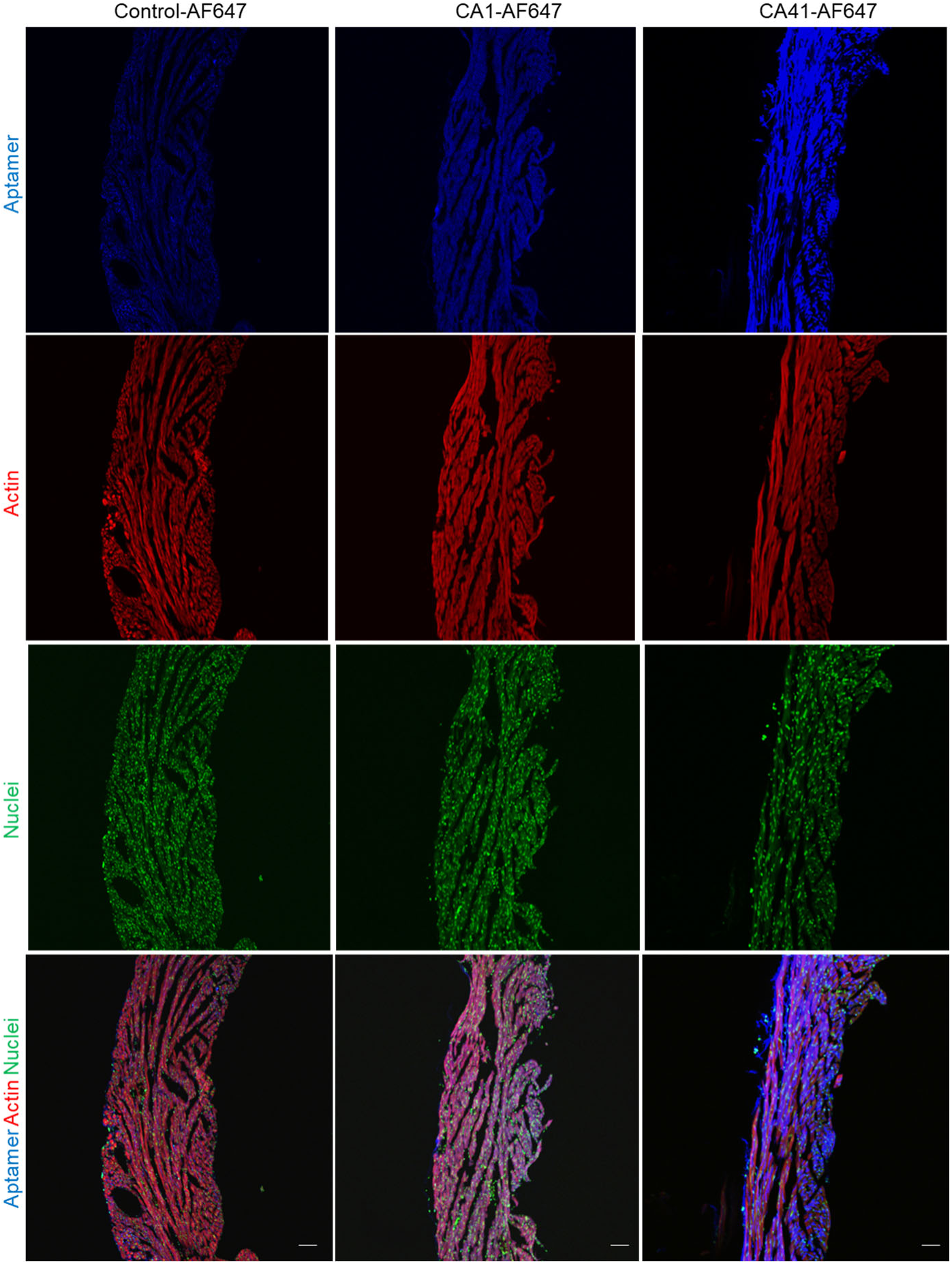
CA1-AF647 targets right ventricle myocardium of mouse hearts perfused *ex vivo*. Aptamer (blue), actin (red), and nuclei (green). Scale = 100 µm

**Supplemental Figure 5:**
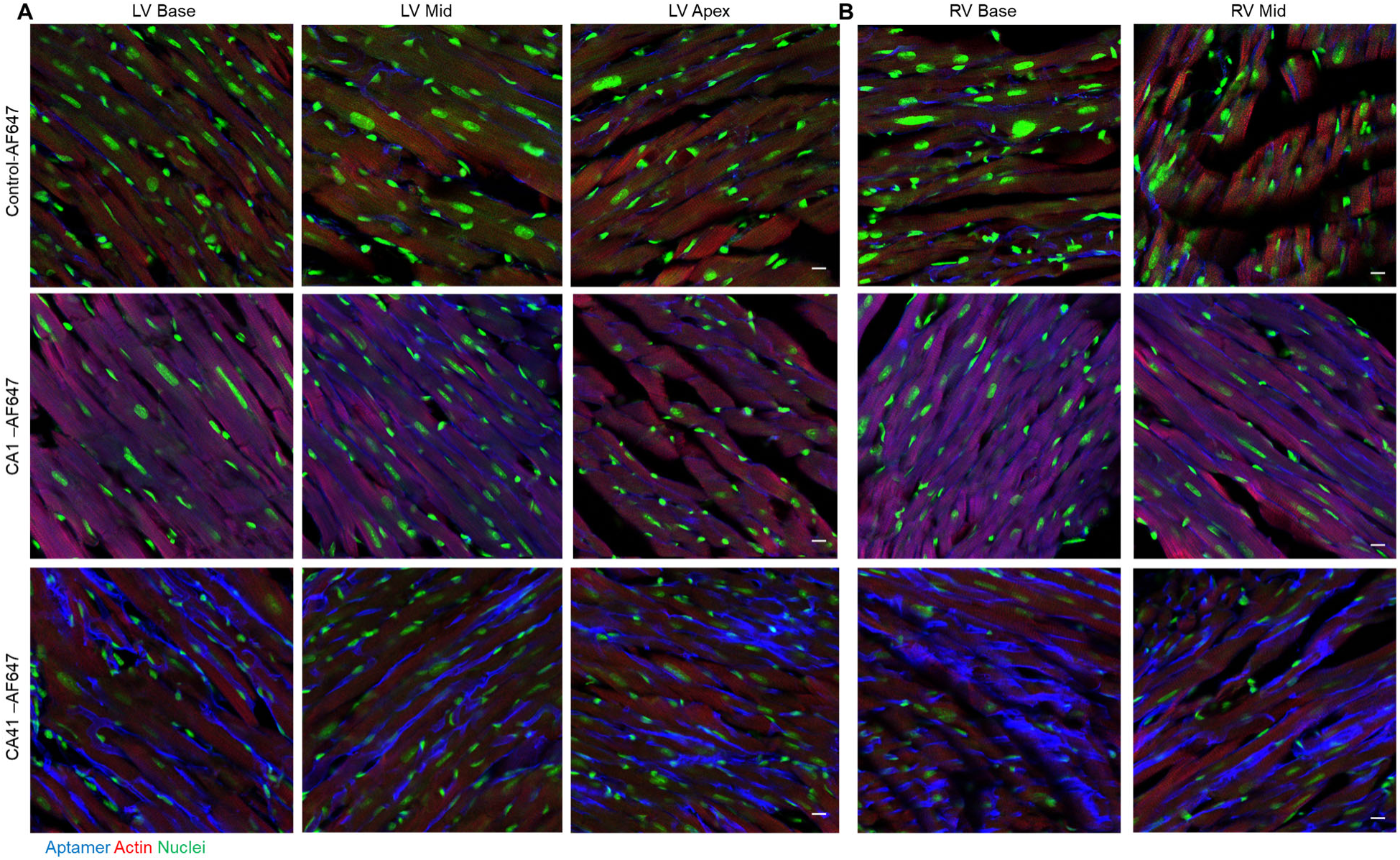
Sections of **A)** left ventricular (LV) and **B)** right ventricular (RV) myocardium of mouse hearts perfused *ex vivo* with either CA1-AF647, CA41-AF647, or control-AF647. Aptamer (blue), actin (red), and nuclei (green). Scale = 25 µm

**Supplemental Table 1**

Redacted

**Supplemental Table 2.**
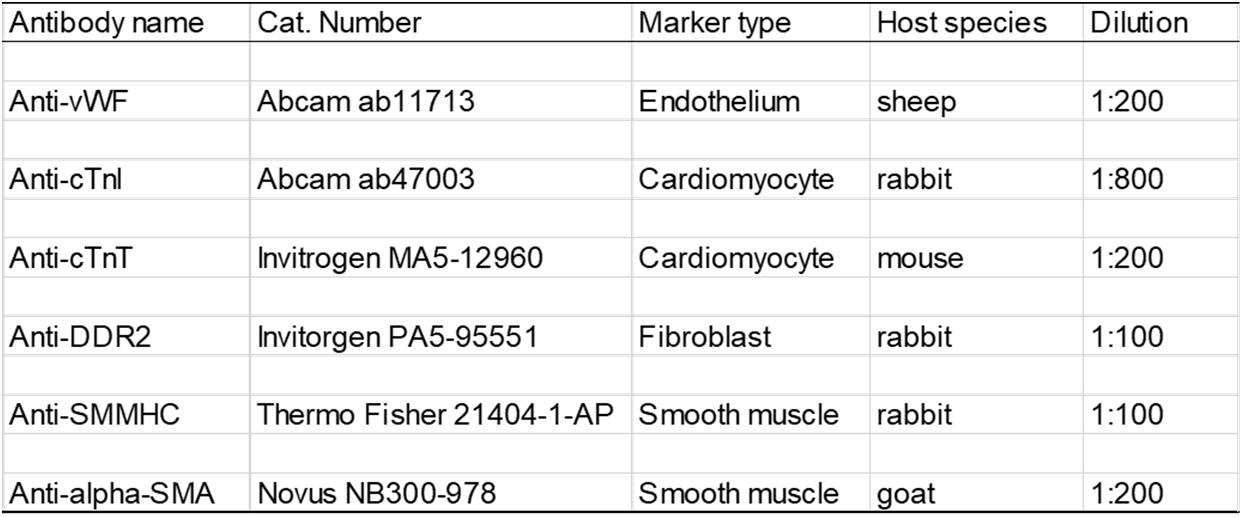

| Antibody name | Cat. Number | Marker type | Host species | Dilution |
| --- | --- | --- | --- | --- |
| Anti-vWF | Abcam ab11713 | Endothelium | sheep | 1:200 |
| Anti-cTnl | Abcam ab47003 | Cardiomyocyte | rabbit | 1:800 |
| Anti-cTnT | Invitrogen MA5-12960 | Cardiomyocyte | mouse | 1:200 |
| Anti-DDR2 | Invitrogen PA5-95551 | Fibroblast | rabbit | 1:100 |
| Anti-SMMHC | Thermo Fisher 21404-1-AP | Smooth muscle | rabbit | 1:100 |
| Anti-alpha-SMA | Novus NB300-978 | Smooth muscle | goat | 1:200 |

